# Repetitive suprathreshold paired-pulse transcranial magnetic stimulation alters corticospinal excitability

**DOI:** 10.64898/2026.10.14.759240

**Authors:** Suraj Suresh, Wei-Yeh Liao, Tahnee Thatcher, Isabelle Villalta, Victoria Prowse, Marissa Holden, Mana Biabani, George M. Opie, Mitchell R. Goldsworthy, Nigel C. Rogasch

## Abstract

**Background:** Suprathreshold paired-pulse transcranial magnetic stimulation (TMS) at interstimulus intervals (ISIs) of 20–30 ms evokes cumulative online increases in corticospinal excitability. Whether these changes persist beyond stimulation remains unclear. This study investigated whether repetitive paired-pulse TMS targeting periods of net excitation or inhibition induces offline changes in corticospinal excitability.

**Methods:** Nineteen healthy participants completed three sessions, separated by at least one week, involving 40 pairs of repetitive paired-pulse TMS delivered to the left primary motor cortex at an ISI of 20 ms (rPPS20) or 100 ms (rPPS100), or a single-pulse control condition (rSPS). Offline changes in corticospinal excitability were assessed using motor evoked potential (MEP) amplitudes recorded before and after stimulation.

**Results:** During stimulation, MEP amplitudes decreased across trials in the rPPS100 condition, although online changes did not differ significantly between conditions. Following stimulation, MEP amplitudes were significantly reduced for up to 40 minutes after rPPS100 (p = 0.017) but were not changed following rPPS20 or rSPS (all p > 0.05).

**Conclusion:** Contrary to our hypothesis, repetitive paired-pulse TMS targeting periods of net excitation did not induce offline changes in corticospinal excitability. In contrast, stimulation targeting long-interval intracortical inhibition (LICI) produced sustained reductions in corticospinal excitability, suggesting that repetitive LICI-inducing stimulation can induce lasting plasticity within the corticospinal system.

## 1. Introduction

Transcranial magnetic stimulation (TMS) is a non-invasive method capable of depolarising cortical neurons across the scalp in humans (Groppa et al., 2012). When delivered over primary motor cortex (M1), TMS activates corticospinal neurons trans-synaptically via indirect (I-wave) connections, leading to a motor evoked potential (MEP) in targeted peripheral muscles (Groppa et al., 2012; Zewdie & Kirton, 2016), the amplitude of which indexes corticospinal excitability (Groppa et al., 2012; Hoogendam et al., 2010). Paired-pulse TMS protocols have additionally been developed to assess inhibitory and excitatory neurotransmission within M1 (Hoogendam et al., 2010). For instance, short-interval intracortical inhibition (SICI) refers to paired-pulse protocols with an interstimulus interval (ISI) of 2-5 ms; these reveal MEP suppression thought to reflect activity of type A gamma aminobutyric acid (GABA-A) receptors (Zewdie & Kirton, 2016; Ziemann et al., 2015). Furthermore, long-interval intracortical inhibition (LICI) refers to MEP suppression with ISIs of 50-200 ms, which is underpinned by GABA type B activation (Ziemann, 2003, 2004; Ziemann et al., 2015). In contrast, intracortical facilitation (ICF) refers to MEP potentiation apparent with ISIs of 10-15 ms, which is thought to reflect processes involving glutamate (Ziemann, 2003, 2004; Ziemann et al., 2015). MEP facilitation can also be observed at ISIs approximating I-wave periodicity (~1.5 ms) (Di Lazzaro et al., 2007; Kidgell et al., 2016; Gary W Thickbroom et al., 2006). These paired-pulse protocols have been used to assess cortical circuits of the primary motor cortex in both healthy brain function and in brain disorders (Hanajima & Ugawa, 2008; Zewdie & Kirton, 2016).

Although paired-pulse TMS has been traditionally used to assess inhibitory and excitatory circuits within M1, several lines of evidence show that these protocols may induce cortical plasticity within targeted circuits that outlasts the period of stimulation. Thickbroom et al. (2006) developed a repetitive paired-pulse TMS protocol targeting a period of net excitation aligned with the first peak of I-wave periodicity (Gary W Thickbroom et al., 2006). This involved 360 pairs of stimuli with an ISI of 1.5 ms and stimulus intensity evoking an MEP with peak-to-peak amplitude of 0.5 mV (G. W. Thickbroom et al., 2006). This intervention produced a 5-fold increase in MEP amplitude during stimulation (i.e., online effects), with MEPs remaining facilitated for 10 mins after stimulation (i.e., offline effects). The authors attributed these changes to increased synaptic efficacy on corticospinal neurons, likely via neural plasticity induced by a long-term potentiation (LTP)-like mechanism. Similar findings have been reported using ISIs targeting other periods of net excitation, including late I-waves (~4.5 ms)(Opie et al., 2021)and ICF (subthreshold conditioning and suprathreshold test stimulus) (Shirota et al., 2016). In contrast, interventions targeting periods of net inhibition, including SICI and LICI, have been mixed, with some reporting a reduction in MEP amplitude, and others reporting no changes (Hoogendam et al., 2010; Shirota et al., 2016). Hence, designing paired-pulse rTMS protocols to target periods of net excitation may represent a reliable strategy to induce synaptic plasticity in humans using TMS (Hanajima & Ugawa, 2008; Kidgell et al., 2016).

In support of this, work from our group recently observed an online increase in MEP amplitudes when delivering suprathreshold paired-pulse TMS (Biabani et al., 2021; Suresh et al., 2026). Specifically, application of 20 stimulus pairs with ISIs of 20 and 30 ms resulted in more than 150% potentiation of MEP amplitude (Suresh et al., 2026). Increases in MEP amplitude across trials were also observed with single-pulse TMS, but not ISIs targeting net inhibition (50-150 ms) (Suresh et al., 2026). However, the single- and paired-pulses were delivered as part of a protocol to assess excitatory and inhibitory circuits, not to induce plasticity (Zewdie & Kirton, 2016). As a result, the experimental protocol randomised 8 paired-pulse conditions with ISIs ranging from 10-200 ms, included single-pulse TMS (200 trials in total separated into 4 blocks), and implemented a jittered inter-trial interval (Suresh et al., 2026). Therefore, it remains unclear whether: 1) similar online increases in MEP amplitude would be achieved if suprathreshold paired-pulse stimuli with one ISI was delivered in a single block with fixed inter-trial interval (as is typical for other repetitive paired-pulse protocols) and 2) whether online effects translate to offline increases in MEP amplitude, indicative of neural plasticity induction.

There were two aims to the current study. First, we assessed whether we could replicate the online (i.e., across trial) increases in corticospinal excitability during single- and paired-pulse TMS when paired stimuli with one ISI were provided as a single block. Second, we assessed whether paired-pulse TMS using suprathreshold conditioning stimuli targeting periods of net excitation or inhibition resulted in offline changes in corticospinal excitability. Based on our recent findings, we hypothesised that corticospinal excitability would increase during application of both single-pulse TMS and paired-pulse TMS with an ISI of 20 ms (targeting net excitation), and these increases would translate to offline potentiation of MEP amplitude. In contrast, we also hypothesised that paired-pulse TMS with an ISI of 100 ms (targeting LICI) would not result in online or offline changes in MEP amplitude

## 2. Methods

### 2.1. Participant

Twenty-one participants were recruited for the study, but only 19 healthy participants (mean age = 31.5 ±2.2 years; range = 18 to 49 years; 11 females, 8 males) successfully completed all experimental sessions. One participant withdrew for personal reasons, whereas we were unable to obtain MEPs in another due to high stimulation threshold. Participants were screened for any contraindications to TMS and provided written informed consent prior to the study. All procedures were approved by the University of Adelaide Human Research Ethics Committee (HREC ID: H-2021-029) in accordance with the National Statement on Ethical Conduct in Human Research from the National Health and Medical Research Council of Australia.

### 2.2 .Experimental overview

Participants attended three experimental sessions that were separated by at least a week to avoid residual inter-session effects. Sessions were conducted at the University of Adelaide’s clinical research facility (CRF). Participants were instructed to keep their eyes open and blink normally whilst seated in a comfortable chair. TMS was used to assess baseline resting motor threshold (RMT) and MEP amplitudes following single-pulse TMS at two baseline time points. Participants then received one of three TMS interventions, the order of which were pseudo-randomised across weeks. These included repetitive paired-pulse TMS with a 20 ms ISI (rPPS20), repetitive paired-pulse TMS with a 100 ms ISI (rPPS100) and repetitive single-pulse TMS (rSPS). MEPs were recorded during the intervention. MEP amplitudes following single-pulse TMS were assessed again at 6 time points up to 40 mins post intervention and compared across conditions (figure 1).

**Figure 1.**
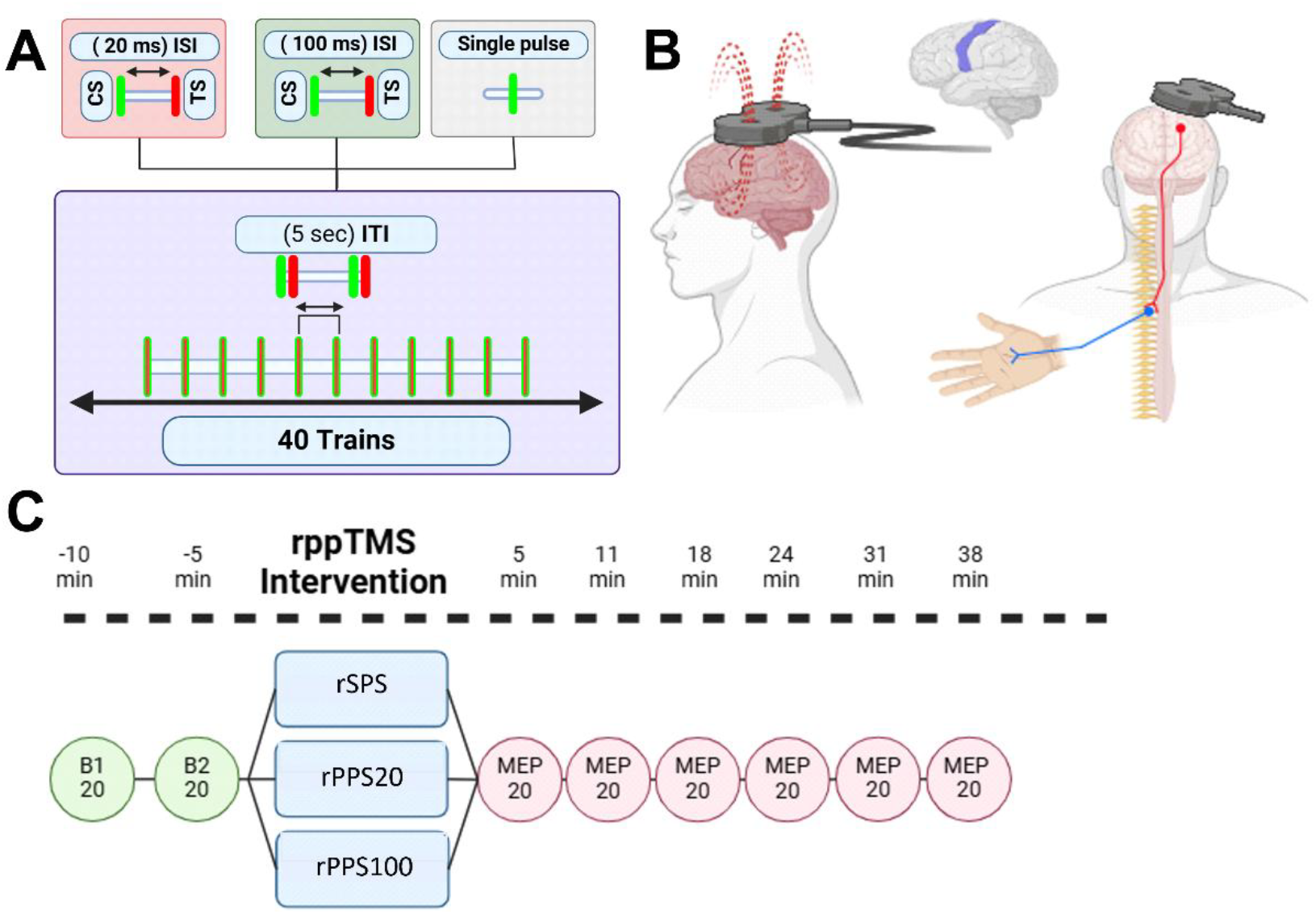
Overview of the experimental design. **A)** Three interventions were tested including: 1) repetitive paired-pulse transcranial magnetic stimulation (rppTMS) with a 20 ms ISI (rPPS20; orange box; ISI = interstimulus interval); 2) rppTMS with a 100 ms ISI (rPPS100; green box); and 3) repetitive single-pulse TMS (rSPS; grey box). The conditioning stimulus (CS) is represented in light green, and the test stimulus (TS) is represented in red. 40 trains of stimulation (e.g., 40 pairs for paired-pulse, 40 single pulses for single-pulse) were given for each intervention with a fixed 5 second inter-train interval (ITI). **B)** Motor-evoked potentials (MEPs) were recorded from the left first-dorsal interosseus muscle using electromyography (EMG) when delivering TMS to the left motor cortex. **C)** Corticospinal excitability was assessed using mean MEPs (n=20 trials) by comparing two timepoints before the intervention (B1, B2) to those taken six timepoints after the intervention starting at 5 mins and spaced by 6.5 mins (5, 11, 18, 24, 31, 38 mins). At each timepoint, TMS was delivered with a 5 second ITI with jittered intervals (±1 second).

### 2.3. EMG

EMG was recorded by applying two Ag-AgCl surface electrodes positioned approximately 2 cm apart over the right first dorsal interosseous (FDI) muscle in a belly-tendon montage. The ground electrode was strapped around the right wrist using a Velcro strap. EMG signals were amplified (×1000), band-pass filtered (10-1000 Hz), digitized at a sampling rate of 5 kHz (CED1902 and CED1401; Cambridge Electronic Design, United Kingdom), and epoched around the TMS pulse (−200 to 500ms).

### 2.4. TMS

Monophasic TMS pulses were administered to M1 using a figure-of-eight coil (D70) connected to two Magstim 200^2^ stimulators (Magstim Company, Whitland, UK) via a BiStim^2^ module. The coil was held tangentially to the scalp with the handle pointing backwards, oriented ~45° to the sagittal plane, resulting in a poster-anterior current direction in the underlying cortex. Throughout the experiment, non-interventional TMS pulses were provided once every 5 seconds with a ± 1 second jitter. At the beginning of each experimental session, the hotspot location was identified as the optimal position over left M1 that consistently produced the largest MEP response in the right FDI muscle. The position was marked on the scalp using a felt-tip pen to enable consistent coil positioning throughout the experiment. RMT was then determined by gradually adjusting the stimulator output to find the minimum intensity that produced MEPs ≥ 50 µV in at least 5 out of 10 trials (expressed as a percentage of maximum stimulator output; %MSO). Stimulation intensity was then increased to find the level that consistently evoked MEPs of 1 mV (S1mV) and 2 mV (S2mV) in peak-to-peak amplitude.

### 2.5. Repetitive paired-pulse TMS interventions

To assess baseline corticospinal excitability, two blocks of single-pulse TMS at S1mV intensity consisting of 20 trials were administered 10 and 5 minutes before the TMS intervention. Next, one of three TMS interventions were pseudo-randomly administered to participants across the three experimental sessions. These interventions included: 1) rPPS20; 2) rPPS100; and 3) rSPS (a control condition). For each intervention, 40 trials were administered (i.e., 80 pulses for paired-pulse conditions, 40 pulses for the single-pulse condition) at a fixed interval of 0.2 Hz.

We chose 40 trials as our previous research indicated increases in corticospinal excitability with ISIs of 20 ms following as few as 20 trials for a given ISI (Suresh et al., 2026). For the paired-pulse conditions, the conditioning stimulus was set to 120% of RMT and the test stimulus to S1mV. For the single-pulse condition, the test stimulus was set to S2mV. We chose a higher stimulation intensity for the single-pulse condition to match the large MEP amplitudes resulting from paired-pulse TMS with a 20 ms ISI (Biabani et al., 2021; Suresh et al., 2026). EMG was recorded throughout the interventions. Following completion of the intervention, 6 blocks of single-pulse TMS at S1mV were repeated at 5, 11, 18, 24, 31, 38 minutes (i.e., 6.5-minute intervals) to assess changes in corticospinal excitability resulting from the interventions.

### 2.6. Data processing

EMG data were processed using Signal software (v4.11, Cambridge Electronic Design). MEP amplitudes were measured by calculating the peak-to-peak amplitude between 20 ms and 40 ms following the test stimulus of each trial. Trials were discarded if EMG activity preceding the MEP exceeded 25 µV.

### 2.7. Statistical analysis

Statistical analysis on TMS data was performed using R Studio (R 4.4.2 + RStudio 202.12.0). TMS intensities (RMT, S1mV, S2mV) obtained at baseline were fit with linear mixed models via the *lme4* (Bates et al., 2015) package investigated the effects of session (rPPS20, rPPS100, rSPS). For MEP amplitude, we fitted generalized linear mixed models with Gamma distribution and log link function using single-trial MEP data, and included random participant intercepts (Lo & Andrews, 2015; Sasaki et al., 2023). A two-factor GLMM assessed the main effects and interactions of session and trial (trials 1–40) on MEP amplitude recorded during the intervention (i.e., online effects). A second two-factor GLMM assessed the fixed effects and interactions of session and timepoint (pre [mean of baseline 1 and 2], and post 5, 11, 18, 24, 31, and 38 minutes timepoints) (i.e., offline effects). Following model fitting, custom pairwise contrasts were generated to probe main effects and interactions via the *emmeans* package (Lenth, 2022). For online effects, we generated estimated trends (and 95% confidence interval; 95% CI) of MEP amplitude across the 40 trials during each intervention and compared trends between sessions (*emtrend* function). For offline effects, we generated estimated marginal means (and 95% CI) and compared MEP amplitude between baseline and post-intervention timepoints, and between interventions at each time point. We also generated baseline-normalised post-intervention MEP amplitude averaged across all timepoints and compared them between interventions. Moreover, linear-regression analysis (Graphpad PRISM) was performed to assess the relationship between online (slope of MEP changes over trials during intervention) and offline (post-intervention MEPs divided by pre intervention MEPs multiplied by 100) changes in corticospinal excitability for each intervention. All comparisons are presented as estimated differences (and 95% CI) for an unstandardised measure of effect size. Bonferroni correction was applied to adjust for type-I errors, with P < 0.05 considered significant.

## 3. Results

### 3.1. Baseline measures

At baseline, there were no differences between sessions for RMT (X^2^_(2,33)_ = 1.76, p = 0.412)), S1mV (X^2^_(2,31)_ = 3.84, p = 0.147) or S2mV (X^2^_(2,32)_ = 3.78, p = 0.151) (Table 1).

**Table 1.**
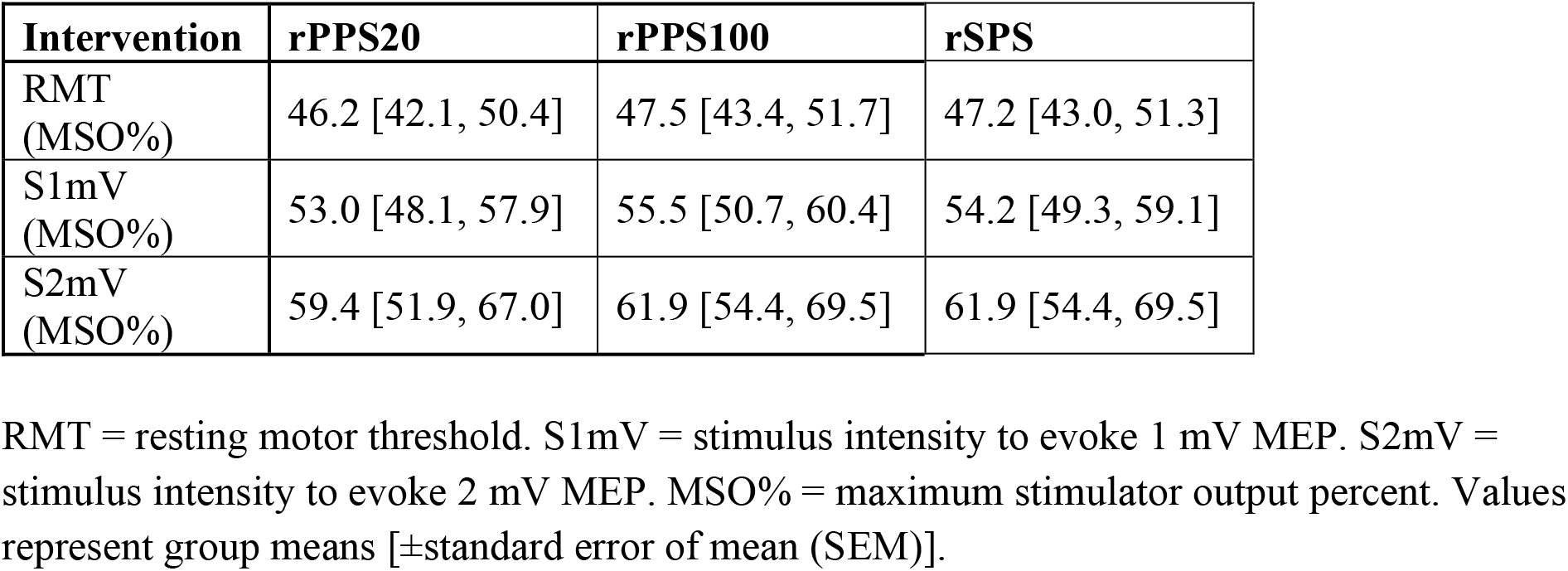
Baseline stimulation intensities in each session.

| Intervention | rPPS20 | rPPS100 | rSPS |
| --- | --- | --- | --- |
| RMT (MSO%) | 46.2 [42.1, 50.4] | 47.5 [43.4, 51.7] | 47.2 [43.0, 51.3] |
| S1mV (MSO%) | 53.0 [48.1, 57.9] | 55.5 [50.7, 60.4] | 54.2 [49.3, 59.1] |
| S2mV (MSO%) | 59.4 [51.9, 67.0] | 61.9 [54.4, 69.5] | 61.9 [54.4, 69.5] |

RMT = resting motor threshold. S1mV = stimulus intensity to evoke 1 mV MEP. S2mV = stimulus intensity to evoke 2 mV MEP. MSO% = maximum stimulator output percent. Values represent group means [ ± standard error of mean (SEM)].

### 3.2. Online effects during TMS interventions

Figure 2 shows the mean amplitude of conditioned MEPs across trials administered during the TMS interventions. There was a significant main effect of intervention (X^2^_(2, 2280)_ = 2315, p < 0.001) and trial (X^2^_(1, 2280)_ = 7.76, p = 0.005), but the intervention x trial interaction was not significant (X^2^_(2, 2280)_ = 0.37, p = 0.831). *Post hoc* analyses assessing the main effect of intervention indicated conditioned MEPs were larger in the rPPS20 condition compared to the rSPS condition (EMD = 37% [21, 54], p < 0.001) and the rPPS100 condition (EMD = 1065% [921, 1229], p = < 0.001). Furthermore, MEPs in the rSPS condition were larger compared to the rPPS100 condition (EMD = 752% [650, 868], p = < 0.001), replicating the well-known effects of paired-pulse TMS at different ISIs on corticospinal excitability.

**Figure 2.**
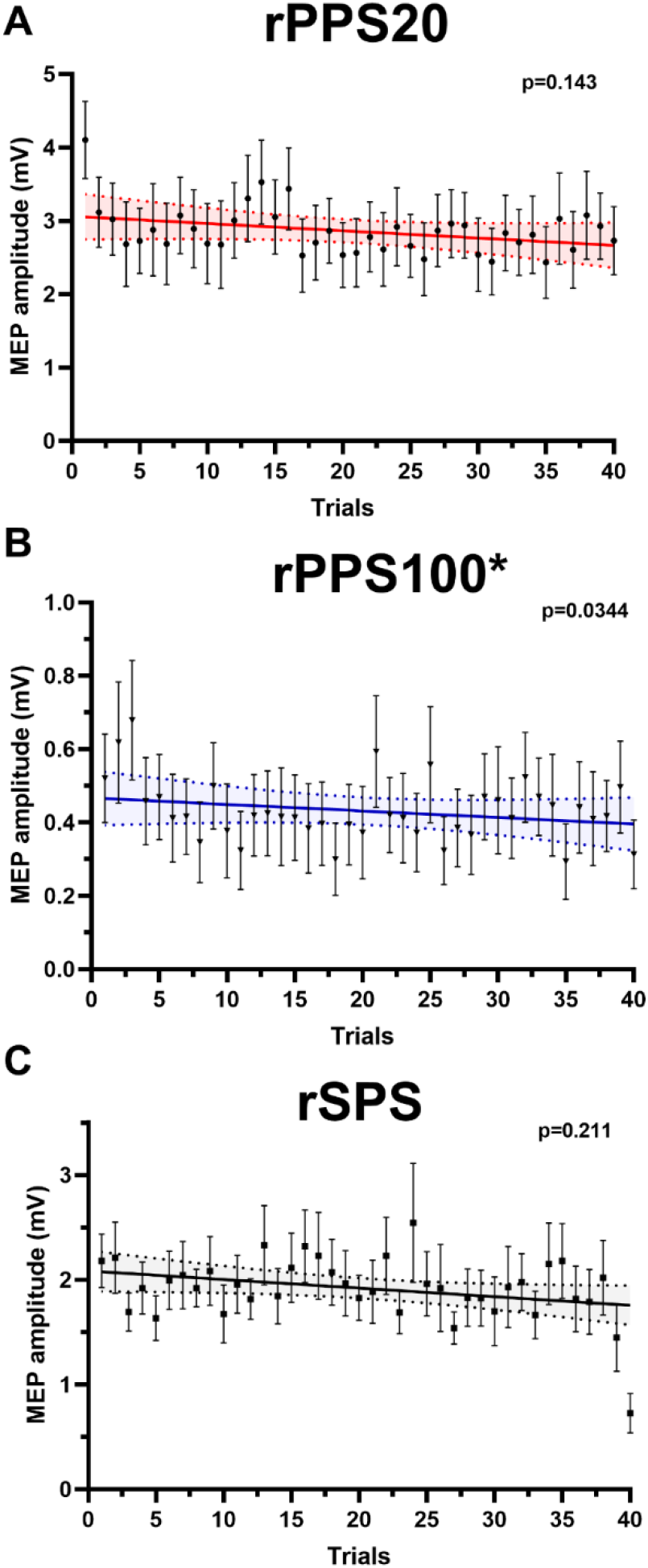
Mean conditioned motor evoked potential (MEP) amplitudes across trials during interventions (online effects). **A)** Repetitive paired-pulse transcranial magnetic stimulation with a 20 ms interstimulus interval (rPPS20). **B)** Repetitive paired-pulse transcranial magnetic stimulation with a 100 ms interstimulus interval (rPPS100). **C)** Repetitive single-pulse transcranial magnetic stimulation (rSPS; control condition). Individual points indicate the mean and standard error (bars) across individuals (n=19) for each trial across the intervention. Horizontal coloured lines indicate the mean line of best fit with confidence intervals (shaded area).

To further investigate the main effect of trial, we compared the slope of the MEP amplitude across trials against ‘0’ (i.e., no change) in each condition. While slopes failed to deviate from a flat line for rPPS20 (estimated trend = −0.004 mV/Trial [−0.01, 0.001], p = 0.143) (Figure 3a) and rSPS (estimated trend = −0.004 mV/Trial [−0.01, −0.002], p = 0.211) (Figure 3b), a significant negative slope was found during rPPS100 (estimated trend = −0.006 mV/Trial [−0.01, −0.0004], p = 0.034; Figure 3c).

**Figure 3.**
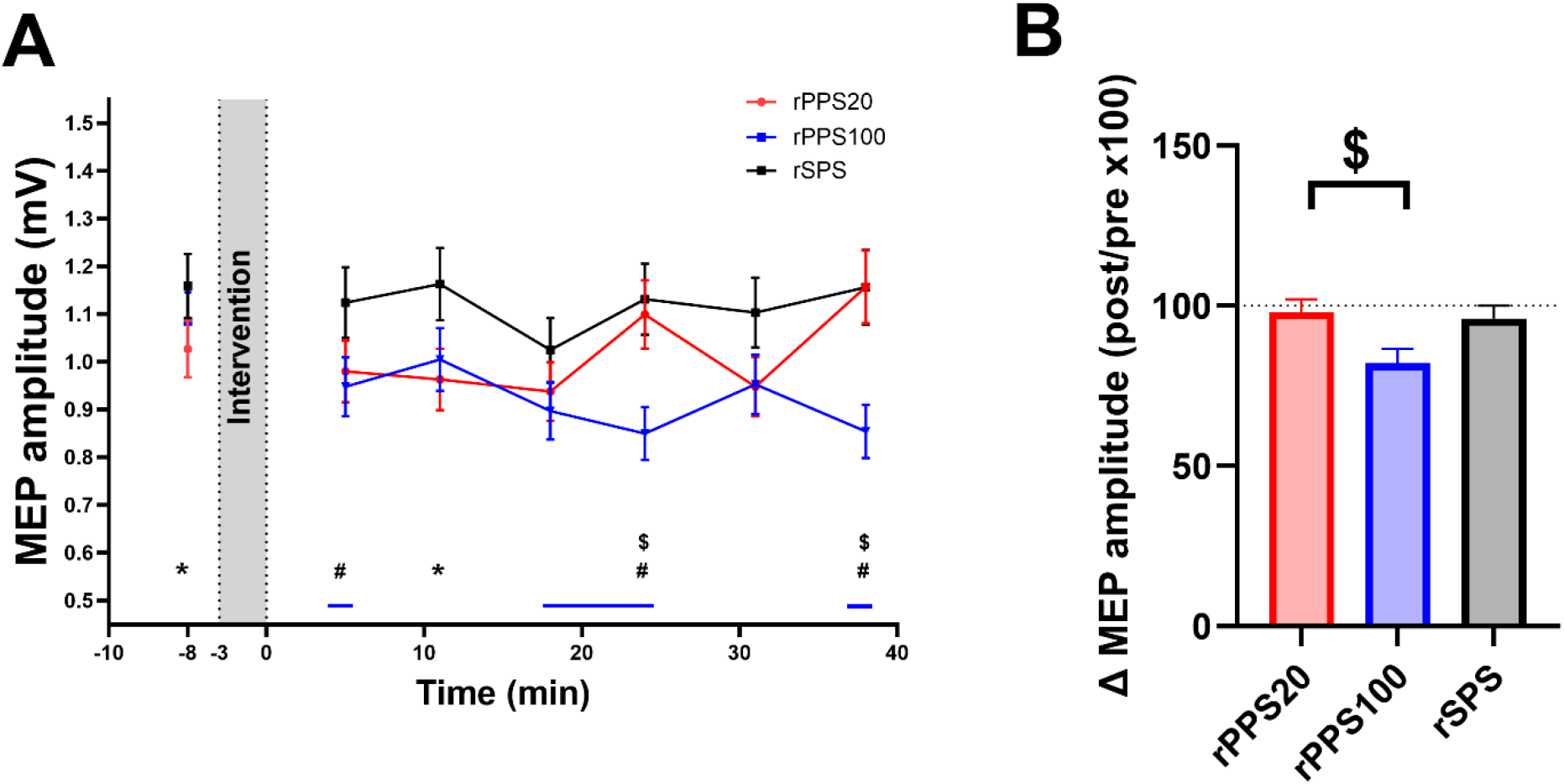
Change in motor evoked potential (MEP) amplitude before and after interventions (offline effects). **A)** Mean (± standard error) MEP amplitudes at each timepoint for each intervention. Note that MEP amplitudes from two baseline timepoints are averaged together. Solid blue lines near x axis indicate timepoints where post MEP amplitudes differed from baseline for the rPPS100 condition (p<0.05). **#** indicates timepoints with significant difference between rPPS100 and rSPS. * indicates timepoints with significant difference between rPPS20 and rSPS. $ indicates timepoints with significant difference between rPPS20 and rPPS100. **B)** Change in MEP amplitudes (mean of post-intervention MEPs divided by mean of pre-intervention MEPs multiplied by 100) following each intervention. * indicates significant difference between conditions (p < 0.05).

### 3.3. Offline effects following TMS interventions

Figure 3 shows the change in MEP amplitude assessed before and after the interventions (i.e., the offline effects). There was a significant main effect of intervention (X^2^_(2, 8485)_ = 50, p < 0.001) and timepoint (X^2^ _(6, 8485)_ = 19, p = 0.004), and a significant intervention x timepoint interaction (X^2^_(12, 8485)_ = 34, p < 0.001) on MEP amplitude. To further assess the intervention x timepoint interaction, we compared MEP amplitudes across timepoints for each intervention, and across interventions at each timepoint (figure 3A). MEP amplitudes were smaller than baseline following rPPS100 at 18 mins (EMD = 17% [1.5, 31], p = 0.021), 24 mins (EMD = 22% [6.7, 34], p < 0.001) and 38 mins (EMD = 21% [6.2, 34], p < 0.001) post intervention. In contrast, responses to rPPS20 and rSPS were not different to baseline at any time post-intervention (all > 0.05). When comparing interventions at different timepoints, MEP amplitudes were smaller in the rPPS100 condition compared to the rSPS condition at the 5 mins (EMD = 16% [1.3, 28], p = 0.028), 24 mins (EMD = 25% [12, 36], p < 0.001), and 38 mins (EMD = 26% [13, 37], p < 0.001) post intervention timepoints, and smaller compared to the rPPS20 condition at the 24 mins (EMD = 23% [9.6, 34], p < 0.001) and 38 mins (EMD = 26% [13, 37], p < 0.001) timepoints. In addition, MEP amplitudes were smaller in the rPPS20 condition compared with the rSPS condition at baseline (EMD = 11% [0.4, 22], p = 0.040) and 11 mins (EMD = 17% [2.9, 30], p = 0.014) post intervention.

To account for potential differences in baseline MEPs between interventions, we also assessed whether the overall change in MEP amplitude from baseline (normalised-MEP amplitude) varied between interventions (figure 3B). Normalised MEP amplitude following rPPS100 were reduced compared to the rPPS20 condition (EMD = 14% [2.1, 25], p = 0.017).

### 3.4. Relationship between online and offline effects and inter-individual variability

Figure 4 shows the relationship between online and offline changes in MEP amplitude, and the inter-individual variability in offline MEP changes for each intervention. A linear regression analysis performed independently for each intervention found no evidence for a relationship between online and offline changes in MEP amplitude (rPPS20, r^2^ = 0.01, p = 0.63; rPPS100, r^2^ = 0.03, p = 0.48; rSPS, r^2^ = 0.11, p =0.16). Furthermore, changes in MEP amplitude in response to each intervention showed substantial inter-individual variability. For rPPS100, only 57% of individuals showed a net decrease in MEP amplitude (43% net increase), despite group-level reductions in MEP amplitude. Individuals showing net decreases/increases were roughly equivalent following both rPPS20 (52/48%) and rSPS (48/52%). Together these findings suggest that online changes in MEP amplitude were not related to offline changes for any intervention, and the individual-level changes in MEP amplitude following all interventions were highly variable.

**Figure 4.**
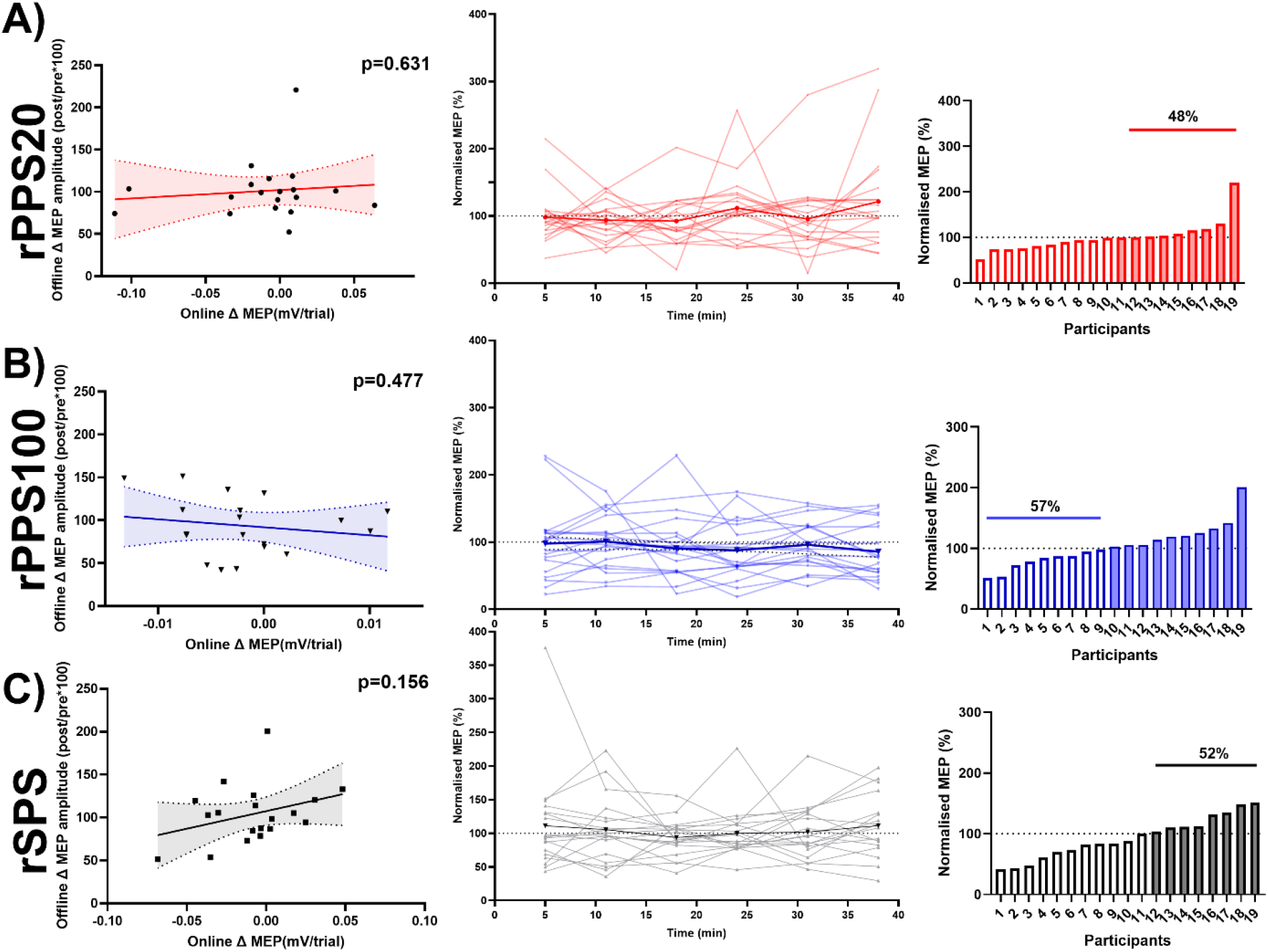
The relationship between online and offline changes in motor-evoked potential (MEP) amplitude, and the inter-individual variability in offline outcomes for each intervention. Left column) Correlation between online and offline changes in MEP amplitude for each intervention. Solid line indicates line of best fit; shaded area represents confidence intervals. Middle column) Changes in MEP amplitude (post/pre*100) across time following each intervention for each individual. Right column) Mean change in MEP amplitude averaged across time following each intervention for each individual. <100% = net decrease in MEP amplitude (white bars); >100% = net increase in MEP amplitude (filled bars). Individuals are ranked from largest decrease to largest increase. Red = rPPS20 (A); blue = rPPS100 (B); black = rSPS (C).

## 4. Discussion

We evaluated whether repetitive suprathreshold paired-pulse stimulation delivered at ISIs targeting net excitation and inhibition altered corticospinal excitability during and after stimulation. Counter to our hypotheses, both rPPS20 and rSPS did not alter corticospinal excitability during and following stimulation. In contrast, rPPS100 produced online reductions in corticospinal excitability that persisted for at least 38 mins after the end of the intervention.

These findings highlight the complex relationship between TMS parameters and neural plasticity, while demonstrating that a brief (40 trial) protocol can modulate corticospinal excitability for over 30 minutes.

### 4.1. Online effects of repetitive paired-pulse TMS

Several studies have shown that rPPS leads to online increases in MEP amplitude. These effects were observed when using ISIs targeting the peaks of I-wave periodicity (ISI = 1.5 ms) (G. W. Thickbroom et al., 2006) and ICF (ISI = 10-15 ms) (Shirota et al., 2016). Similar cumulative MEP increases have also been observed during trains of single-pulse TMS, particularly using inter-trial intervals < 10 s (Julkunen et al., 2012; Pellicciari et al., 2016; Suraj Suresh, Unpublished)). In a recent study by our group, we noted cumulative online increases in MEP amplitude when using a paired-pulse protocol designed to assess excitatory and inhibitory circuits (8 ISIs between 10-200 ms randomised across trials, suprathreshold conditioning pulse, 200 trials in total) (Julkunen et al., 2012; Suresh et al., 2026). Despite only 20 trials for each ISI, increases were apparent for ISIs targeting periods of net excitation (20 and 30 ms), in addition to following single pulses, but were absent at longer ISIs associated with LICI (50-150 ms) (Raco et al., 2017; Suresh et al., 2026). Furthermore, the findings replicated across two independent cohorts, suggesting that a short stimulation protocol targeting excitatory circuits at 20 or 30 ms ISI may effectively increase M1 excitability. Based on these findings, we expected a short block (e.g., 40 trials) of rPPS20 to show online increases in MEP amplitude, as would the rSPS condition. Contrary to our hypothesis, we found both rPPS20 and rSPS did not modulate MEP amplitude over the course of the stimulation period, whereas MEPs reduced during rPPS100.

As we have previously observed an increase in MEP amplitude online with rPPS20 within two independent cohorts(Biabani et al., 2021; Suresh et al., 2026), it is unclear why we did not replicate this effect in the present study. While sources of participant variability may have played a role, including attention, cortisol levels, genetics, physical activity and chronotype (Ridding & Ziemann, 2010) it is likely that the rPPS20 stimulation parameters were not effective in the present study. Notably, our previous assessments of paired-pulse TMS nested eight ISIs and single-pulse TMS within a single-block, which was repeated four times and totalled 200 trials (Biabani et al., 2021; Suresh et al., 2026). While we observed specific online MEP modulation at 20 and 30 ms ISIs in these earlier studies, it is possible that these cumulative increases in MEP amplitudes were the result of complex interactions between different excitatory and inhibitory circuits that were activated by single and multiple paired-pulse TMS conditions occurring over a larger number of trials. Regardless, our findings suggest that specifically targeting excitatory circuits over a shorter period of stimulation may not be effective in modulating cortical excitability. In contrast, targeting inhibitory circuits at 100 ms ISI in isolation appears to be effective and leads to a decrease in MEP amplitude. This new finding suggests that changes in excitatory and inhibitory circuits over the course of a stimulation block may reflect different mechanisms. For example, potentiation of excitatory circuits may require a longer block of stimulation or concurrent activation of other cortical circuits. In contrast, inhibitory circuits can be modulated within a shorter period of stimulation without the activation of other neural circuits. While the mechanisms of rPPS100 will need to be explored in future studies, it may be a useful alternative for reducing cortical excitability online.

### 4.2. Offline effects of repetitive paired-pulse TMS

Contrary to our hypotheses, the post-intervention assessment found that only the rPPS100 condition altered corticospinal excitability beyond the period of stimulation, with MEP amplitudes decreasing for up to 38 minutes. These changes in corticospinal excitability differed from those evoked by rPPS20, but not the rSPS condition. Reductions in MEP amplitude associated with the rppTMS100 condition may stem from neuroplastic reductions in synaptic communication, possibly involving LTD-like mechanisms. Alternatively, the intervention may have increased GABAergic neurotransmission within the cortical circuits underlying LICI, which are primarily mediated by the GABA_b_ receptor (Darmani & Ziemann, 2019; McDonnell et al., 2006; Ziemann, 2003; Ziemann et al., 2015). Our findings differ from those reported by Shirota et al. (2016), who reported that rPPS targeting ICF circuits (~ISI of 15 ms; subthreshold conditioning pulse) produced increases in corticospinal excitability lasting 20 mins following the intervention, whilst rPPS targeting LICI circuits (~ISI of 100 ms; suprathreshold conditioning pulse) produced no change in corticospinal excitability. Although intervention parameters, such as stimulation intensity and ISI, may explain some of the inconsistencies between the studies, the most notable difference was the number of stimulation pairs included in the intervention. Shirota and colleagues used 360 pairs of stimuli (30 mins), whereas we used 40 pairs of stimuli (3 mins 20 s) (Shirota et al., 2016). We chose a relatively short intervention period based on our previous online findings, which showed online changes in MEP amplitude across 20 trials for 20 and 30 ms ISI pairs (although this was part of 200 trials in total) (Suresh et al., 2026).

The duration of stimulation appears to play a key role in determining the direction of plasticity following rTMS. For example, for theta burst stimulation, the direction of plasticity differs with 600 pulses compared to either 300 or 1200 pulses (Gamboa et al., 2010; Gentner et al., 2008; Nettekoven et al., 2014). Modelling studies have suggested this variability may result from attractor dynamics emerging from the molecular mechanisms governing meta-plasticity, where the direction of plasticity oscillates during stimulation regardless of the protocol parameters (Fung & Robinson, 2014; Repo, 2024; Thomson & Sack, 2020; Wilson et al., 2018; Wilson et al., 2021). While speculative, it is possible that similar mechanisms may also impact plasticity outcomes following rPPS protocols (Huang et al., 2007; Nakazono et al., 2021). Future work assessing the impact of different train lengths on rPPS outcomes is required to further clarify this possibility.

### 4.3. Implications

These findings illustrate the complex relationship between TMS parameters and corticospinal excitability modulation. First, while earlier studies demonstrated a relationship between online and offline modulation of M1 excitability with rPPS (Kimura et al., 2013; Meadows et al., 2026; Sewerin et al., 2011; G. W. Thickbroom et al., 2006), our results are more consistent with recent work suggesting that online and offline effects may be mechanistically different (Sasaki et al., 2024; Meadows et al., 2026). Second, the findings suggest that single and paired-pulse paradigms designed to assess neurophysiological processes can modulate corticospinal excitability over the course of the stimulation block, in contrast to the traditional view that these paradigms are not expected to influence neural circuits. Notably, the rPPS100 condition resulted in cumulative online changes in corticospinal excitability. This suggests that researchers will need to be cautious when assessing LICI during longer blocks of stimulation.

Furthermore, we showed that a paired-pulse protocol targeting these inhibitory circuits resulted in a reduction in corticospinal excitability that persisted up to 40 mins following the intervention. Longer LICI blocks may therefore influence subsequent neurophysiological measures of cortical excitability (Daskalakis et al., 2008; Opie et al., 2017; Opie et al., 2018; Premoli et al., 2014; Rogasch et al., 2013; Tremblay et al., 2019). Despite these considerations, our findings also suggest that rPPS100 could be an effective protocol for reducing corticospinal excitability, complementing current rPPS protocols that increase excitability. Future research will need to assess the intervention dosing and compare rPPS100 with other protocols designed to decrease corticospinal excitability like 1 Hz rTMS and continuous theta burst stimulation.

### 4.4. Limitations

Certain limitations should be considered when interpreting the results of the current study. First, modulations in both cortical and/or spinal excitability can influence MEP amplitude, making it difficult to determine at which level changes in excitability are occurring during and after TMS protocols (Di Lazzaro et al., 2018; Kujirai et al., 1993; Nakamura et al., 1997; Valls-Solé et al., 1992). TMS combined with EEG is an alternative method which more directly reflects the excitability of cortical circuits (Rogasch & Fitzgerald, 2013), whereas methods like the H-reflex and F-waves can provide information on spinal excitability (Inghilleri et al., 2003). Combining these methods could provide a more detailed understanding of where excitability changes are taking place during and after rPPS. Second, the number of trains delivered in this study (40 trains) is substantially fewer than the 180-360 trains typically administered in previous rPPS studies. While we deliberately chose a small number of trains based on our previous findings, it is possible that we simply did not apply a long enough protocol to evoke increases in corticospinal excitability.

### 4.5. Conclusion

The study suggests that rPPS20 and rSPS are not effective stimulation modalities to evoke lasting changes in corticospinal excitability. However, rPPS100 reduced corticospinal excitability for up to 40 mins and only required 40 trains of stimulation (just over 3 minutes). As such, rPPS100 represents a potential new protocol for rapidly reducing corticospinal excitability.

## Author contributions

Suraj Suresh: Conceptualization, Methodology, Formal analysis, Investigation, Data Curation, Writing - Original Draft, Visualization.

Wei-Yeh Liao: Formal analysis, Writing - Review & Editing. Tahnee Thatcher: Investigation, Writing - Review & Editing. Isabelle Villalta: Investigation, Writing - Review & Editing. Victoria Prowse: Investigation, Writing - Review & Editing. Marissa Holden: Investigation, Writing - Review & Editing.

Mana Biabani: Conceptualization, Methodology, Supervision, Writing - Review & Editing. George M. Opie: Conceptualization, Methodology, Supervision, Writing - Review & Editing. Mitchell R. Goldsworthy: Conceptualization, Methodology, Project administration, Supervision, Writing - Review & Editing, Funding acquisition.

Nigel C. Rogasch: Conceptualization, Methodology, Project administration, Supervision, Writing - Review & Editing, Funding acquisition.

## Declaration of generative AI use

During the preparation of this work the authors used ChatGPT in order to assist with language editing, improving clarity and structure of the manuscript. After using this tool/service, the authors reviewed and edited the content as needed and take full responsibility for the content of the published article.

